# High number of HPAI H5 Virus Infections and Antibodies in Wild Carnivores in the Netherlands, 2020-2022

**DOI:** 10.1101/2023.05.12.540493

**Authors:** Irina V. Chestakova, Anne van der Linden, Beatriz Bellido Martin, Valentina Caliendo, Oanh Vuong, Sanne Thewessen, Tijmen Hartung, Theo Bestebroer, Jasja Dekker, Bob Jonge Poerink, Andrea Gröne, Marion Koopmans, Ron Fouchier, Judith M.A. van den Brand, Reina S. Sikkema

**Author notes:** Corresponding author: Reina S. Sikkema, ErasmusMC, department Viroscience, Wytemaweg 80, 3015 CN Rotterdam, The Netherlands. contributed equally (Irina V. Chestakova, Anne van der Linden, Judith M.A. van den Brand, Reina S. Sikkema).

## Abstract

In October 2020, a new lineage of clade 2.3.4.4b HPAI virus of the H5 subtype emerged in Europe, resulting in the largest global outbreak of HPAI to date, with unprecedented mortality in wild birds and poultry. The virus appears to have become enzootic in birds, continuously yielding novel HPAI virus variants. The recently increased abundance of infected birds worldwide increases the probability of bird-mammal contact, particularly in wild carnivores. Here, we performed molecular and serological screening of over 500 dead wild carnivores for H5 HPAI virus infection and sequencing of positive materials. We show virological evidence for HPAI H5 virus infection in 0.8%, 1.4% and 9.9% of animals tested in 2020, 2021 and 2022 respectively, with the highest proportion of positives in foxes, polecats and stone martens. We obtained near full genome sequences for seven viruses and detected PB2 amino acid substitutions known to play a role in mammalian adaptation in three of these. Infections were also found in animals without associated neurological signs or mortality. Serological evidence for infection was detected in 20% of the study population. These findings suggest that a higher number of wild carnivores are infected but undetected in current surveillance programs. We recommend increased surveillance in susceptible mammals, irrespective of the presence of neurological signs or encephalitis.

## Introduction

Wild birds, particularly those belonging to the orders Anseriformes and Charadriiformes, are the natural host of a wide range of low pathogenic avian influenza (LPAI) viruses. In poultry, viruses of the H5 and H7 subtypes can evolve into highly pathogenic avian influenza (HPAI) viruses, which can cause severe disease and mortality in domestic and wild birds. After the emergence of the HPAI A/Goose/Guangdong/1/96 (GsGd) lineage in China, HPAI virus infection was frequently detected in wild birds, causing significant mortality in some species [1,2]. After 2004, descendants of the GsGd H5 viruses spread to Europe via infected migratory birds, and caused global outbreaks in poultry and wild birds[2] In October 2020, a new lineage of clade 2.3.4.4b HPAI H5 virus emerged in Europe and subsequently spread to the Americas [3,4]. This resulted in the largest global outbreak of HPAI so far, with unprecedented mortality in wild birds as well as poultry. Moreover, the epidemiology of HPAI H5 virus seems to have shifted, with enzootic circulation, leading to year-round virus presence and local generation of novel reassortants [3,5].

Mammal infections with HPAI H5 viruses have been described previously [6], but less frequent as compared to the current HPAI H5 global outbreak. This includes reports of infections in wild and domestic carnivores [7-9] as well as sea mammals [10] [11] [12]. In some cases, mutations that have been associated with adaptation to replication in mammals were described [8]. To date, there is no definitive evidence of transmission amongst wild mammals. However, the recent HPAI H5N1 virus outbreak in a mink farm in Spain seemed to indicate that mammal-to-mammal transmission is possible [13]. Large outbreaks in seals (*Phoca vitulina; Halichoerus grypus*) in the U.S. [14] and mass mortality of South American sea lions (*Otaria flavescens)* in Peru may also point to spread in mammal populations [15]. The frequent spillovers and the widespread infections, including some evidence for mammal to mammal transmission, raise concerns on the possibility of further adaptation to mammals [13,16].

In order to study the impact of the current HPAI H5 epizootic on wild carnivores, with possible consequences for public health, wild carnivores were collected and tested for HPAI H5 virus in the period 2020-2022. Included animals were either dead or diseased wild carnivores reported by the general public, or raccoons and stone martens that were trapped and subsequently euthanized outside of this project. Moreover, included carnivores were subjected to antibody testing. Combining molecular, serological and sequence information will contribute to increased knowledge on disease presentation, incidence and possible adaptation of HPAI H5 viruses in wild carnivores. This will provide valuable input for One Health risk analyses and prevention and control measures.

## Material and Methods

### Animal sample collection

Wild carnivores (n=188) that were reported dead or ill by the public as part of a citizen reporting system of dead wildlife were submitted to the Dutch Wildlife Health Centre (DWHC) between 2020-2022. Included species are: Pine marten (*Martes martes*), Polecat (*Mustela putorius*), Badger (*Meles meles*), Stoat (*Mustela erminea*), Marten (species undetermined; *Martes spp*.), Otter (*Lutra lutra*), Raccoon (*Procyon lotor*), Stone marten (*Martes foina)*, Fox (*Vulpes vulpes*), Weasel (*Mustela nivalis*) and Wolf (*Canis lupus)*. Some raccoons had been euthanized as part of the national invasive animal control policy. Additionally, stone martens (n=375) were collected and frozen after they were euthanized in a pilot program to study the effect of culling stone martens on the breeding success of meadow birds in agricultural areas [17]. Stone martens were sampled in farmland across the province of Friesland, in the north of the Netherlands. In addition, 73 serum samples collected by hunters from foxes in the Netherlands in 2017 in a separate study (https://dwhc.nl/vossenonderzoek-kjv/, retrieved on 14 March 2023) were included, as well as blood samples collected from one polecat (2017) and three stone martens (2016) admitted to the DWHC.

In the Netherlands there are estimated to be 5500 badgers [18]; around 100.000 stone martens [19]; 450 otters [20]; between 111.000 and 222.000 red foxes [21] and at least 18 wolves (https://www.bij12.nl/wp-content/uploads/2022/12/Tussenrapportage-wolf-21-december-2022.pdf). For the other included species, no published population estimates are available.

Oropharyngeal and rectal swabs were taken in 1.2 ml virus transport medium for virological testing and stored at -80°C before processing. Lung and brain samples were taken during necropsy and stored at -80°C before processing. Blood (clots) or body fluid was collected for serology and stored at -80°C until use. Due to severe autolysis or severe trauma, not all samples could be taken from all animals

### Sample processing

When available, oropharyngeal and rectal swabs and lung and brain tissues of each individual animal were processed and tested. All samples were placed in lysis buffer and screened under biosafety level 2 conditions. HPAI H5 virus positive samples were handled under biosafety level 3 conditions, e.g. for virus isolation attempts in cell culture. Swab and lung samples were processed individually between June 2020 and May 2021. From June 2021 until end of 2022, both sample types were pooled. A small fraction of about 4×4 mm from each tissue was transferred to a 2 mL vial with a 1/4” ceramic sphere (MP Biomedicals, Solon OH, USA) and 300 μL MagNA Pure 96 DNA Tissue Lysis Buffer (Roche Diagnostics GmbH, Mannheim, Germany). Homogenization took place using the FastPrep-24 5G Homogenizer (MP Biomedicals). Homogenates were cleared by centrifugation at 17,000 xg during 5 minutes and the supernatants were diluted 1:10 in Virus Transport (VTM) medium. Blood samples obtained from culled animals were partially clotted in the tube and the serum was used for testing.

### RNA extraction

From June 2020 – January 2021, RNA/DNA extraction from swab material was performed by an in-house method of manual extraction using magnetic beads, as described previously [22]. RNA/DNA extraction from tissue material was performed on a MagNA Pure LC instrument, with 600 μL of the diluted supernatant and 600 μL MagNA Pure 96 External Lysis Buffer (Roche LifeScience, Basel, Switzerland) as input. From February 2021 onwards, RNA/DNA extraction from all material types was performed on the MagNA Pure 96 platform (Roche Diagnostics GmbH, Mannheim, Germany). Phocine Distemper Virus (PDV) was added and used as internal control as described previously [23].

### Real-time RT-PCR

All samples were tested for the influenza A virus matrix gene [24], but with updated primers (see below) and combined in a duplex reaction with PDV [25]. All influenza A matrix positive samples were subsequently tested in a H5-specific PCR reaction designed in house[24], with updated primers. From November 2022 onwards, samples were tested in a triplex reaction for the influenza A virus matrix gene combined with the H5 gene and PDV as internal control. Fit point analysis was used to determine the Ct values and the cut-off threshold was set manually above the background signals of the negative controls. Samples with a Ct above 40 were considered negative. Updated matrix primers and probes were: 5′ CTT CTR ACC GAG GTC GAA ACG TA 3′ (forward), 5′TCT TGT CTT TAG CCA YTC CAT GAG 3′ (reverse), 5′ FAM-TCA GGC CCC CTC AAA GCC GAG A-BHQ1 3′ (probe 1) and 5′ FAM-TCA GGC CCC CTC AAA GCC GAA A-BHQ1 3 (probe 2). Updated H5 primers and probes were: 5′ GAG AGG AAA TAA GTG GAG TAA AAT TGG A 3′ (forward), 5′ AAG ATA GAC CAG CTA CCA TGA TTG C 3′ (reverse), 5′ FAM-TTT ATT CAA CAG TGG CGA GTT CCC TAG CAC T-TAMRA 3′ (probe used until September 2022), 5′ YakimaYellow-TTT ATT CAA CAG TGG CGA GTT CCC TAG CAC T-BHQ1 3′ (probe used since October 2022).

### Multi segment RT-PCR and whole-genome sequencing

To determine the whole genome consensus sequence of HPAI H5 viruses, RNA was re-extracted from original material using the High Pure RNA Isolation Kit (Roche Diagnostics GmbH Mannheim). A multi segment RT-PCR amplification was performed using the Superscript III high-fidelity RT-PCR Kit (Invitrogen, USA). Influenza virus specific primers were used, containing 13 conserved nucleotides at the 5′terminus and 12 nucleotides and unique barcoded primers at the 3′terminus, covering all eight Influenza segments [26].

The libraries were generated using a ligation sequencing kit (SQK-LSK109, Oxford Nanopore technologies) and multiplexed and sequenced on a MinION R9 flowcell (Oxford Nanopore technologies) according to the manufacturer’s instructions. Porechop software was used to demultiplex the reads that contained a barcode. For analysis, FASTQ-files were imported to the CLC Genomics Workbench v20.0.03 (QIAGEN) and analyzed as described previously [22].

Sequences were mapped to the following reference sequences (GISAID ID):*EPI_ISL_5804788 A/Mute Swan/Netherlands/21037283-002/2021; EPI_ISL_890664 A/Mallard Duck/Netherlands/32/2011 LPAI H5N2; EPI_ISL_890124_A/Mallard/Netherlands/32/2011 NA H5N2; EPI_ISL_1841835_A/Duck/Netherlands/18018989-011015/2018 NA H5N3; EPI_ISL_1774274_A/Anas_Platyrhynchos/Belgium/10811_6/2019 NA H5N6_NA; EPI_ISL_1839504_A/turkey/England/038730/2020_NA_H5N8_NA*. To extract the consensus genomes the following parameters were used: match score=1, mismatch cost=2, length fraction=0.7 and similarity fraction=0.8.

Sequences were submitted to GISAID [27]: EPI_ISL_12066188, EPI_ISL_12069288, EPI_ISL_12069289, EPI_ISL_13201074, EPI_ISL_14393097, EPI_ISL_17583227 and EPI_ISL_17583228.

### Protein microarray (PMA)

Nitrocellulose glass slides with influenza virus antigens were prepared as described previously [28-31]. In short, commercially available NP and HA proteins (Sino Biological, Eschborn, Germany and Immune Technology, New York, USA) were mixed with protein array buffer (Maine manufacturing, GVS Group, Italy) including 4 μl/ml EZ block™ protease inhibitor cocktail (Bio Vision, Waltham, USA). Proteins were spotted in duplicate onto a nitrocellulose 64 pad coated UniSart glass slide (Sartorius Stedim Biotech, Goettingen, Germany) or 24 pad coated AVID glass slide (Grace Bio-Labs, Bend, USA) using a non-contact sci-Flex array spotter (Scienion, Berlin, Germany). Concentrations were determined as described previously [28].

All samples were screened using the nucleoproteins of an H7N9 virus A/Anhui/1-BALF_RG6/2013 (Sino Biological) and H1N1 virus A/California/07/2009 (Sino Biological) using an arbitrary fluorescence cut-off of 6,000. When sufficient material was left, blood samples above cutoff were subsequently tested for HA1 binding antibodies against hemagglutinin subtypes H1-H9, H11 and H16 (table 1). The cut-off for HA1 was 40.000 PFU (dilution 1:32), based on validation by Freidl et al [30].

**Table 1.**
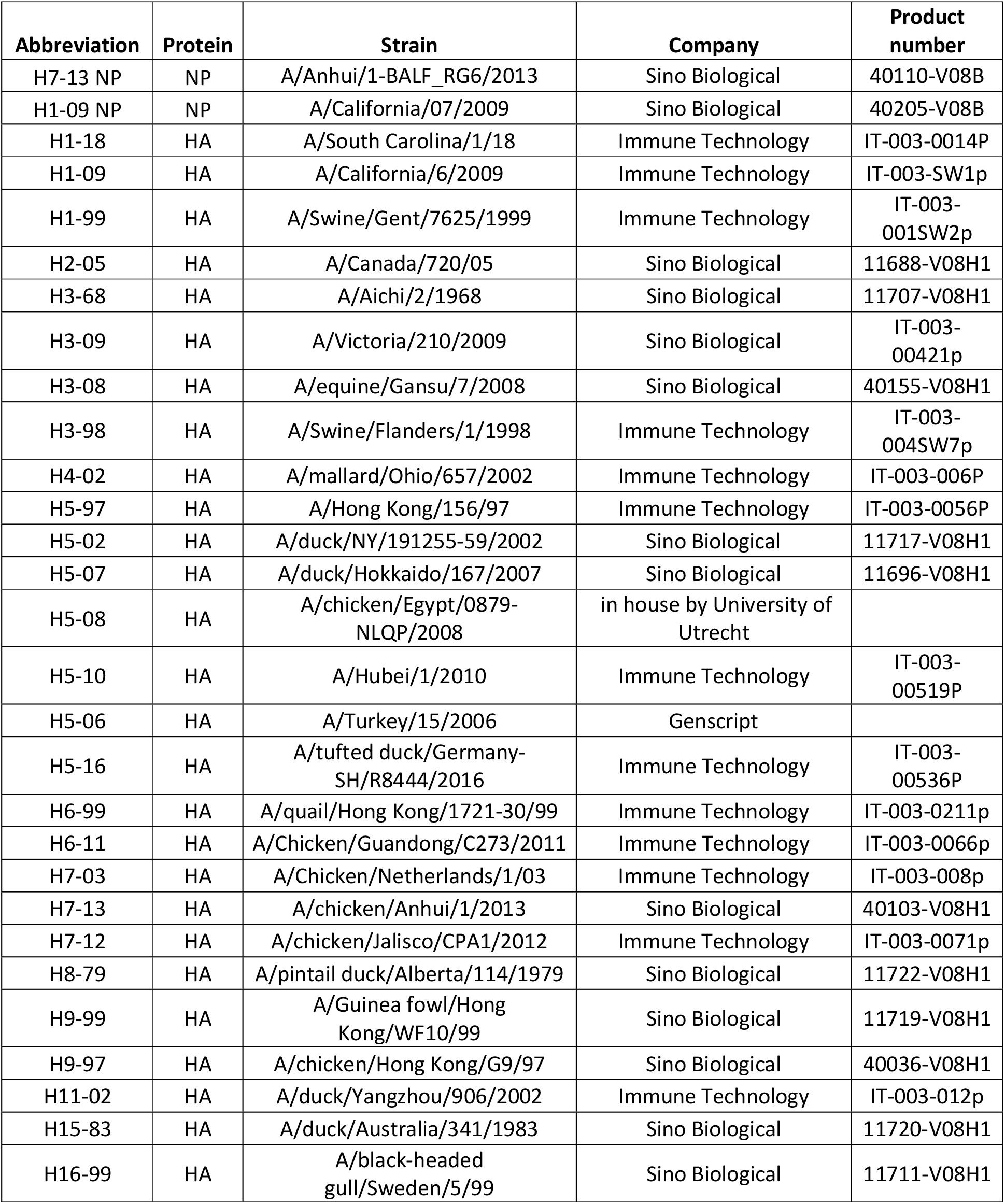
Antigens (NP and HA1) spotted on protein array slides

| Abbreviation | Protein | Strain | Company | Product number |
| --- | --- | --- | --- | --- |
| H7-13 NP | NP | A/Anhui/1-BALF_RG6/2013 | Sino Biological | 40110-V08B |
| H1-09 NP | NP | A/California/07/2009 | Sino Biological | 40205-V08B |
| H1-18 | HA | A/South Carolina/1/18 | Immune Technology | IT-003-0014P |
| H1-09 | HA | A/California/6/2009 | Immune Technology | IT-003-SW1p |
| H1-99 | HA | A/Swine/Gent/7625/1999 | Immune Technology | IT-003-001SW2p |
| H2-05 | HA | A/Canada/720/05 | Sino Biological | 11688-V08H1 |
| H3-68 | HA | A/Aichi/2/1968 | Sino Biological | 11707-V08H1 |
| H3-09 | HA | A/Victoria/210/2009 | Sino Biological | IT-003-00421p |
| H3-08 | HA | A/equine/Gansu/7/2008 | Sino Biological | 40155-V08H1 |
| H3-98 | HA | A/Swine/Flanders/1/1998 | Immune Technology | IT-003-004SW7p |
| H4-02 | HA | A/mallard/Ohio/657/2002 | Immune Technology | IT-003-006P |
| H5-97 | HA | A/Hong Kong/156/97 | Immune Technology | IT-003-0056P |
| H5-02 | HA | A/duck/NY/191255-59/2002 | Sino Biological | 11717-V08H1 |
| H5-07 | HA | A/duck/Hokkaido/167/2007 | Sino Biological | 11696-V08H1 |
| H5-08 | HA | A/chicken/Egypt/0879-NLQP/2008 | in house by University of Utrecht |  |
| H5-10 | HA | A/Hubei/1/2010 | Immune Technology | IT-003-00519P |
| H5-06 | HA | A/Turkey/15/2006 | Genscript |  |
| H5-16 | HA | A/tufted duck/Germany-SH/R8444/2016 | Immune Technology | IT-003-00536P |
| H6-99 | HA | A/quail/Hong Kong/1721-30/99 | Immune Technology | IT-003-0211p |
| H6-11 | HA | A/Chicken/Guandong/C273/2011 | Immune Technology | IT-003-0066p |
| H7-03 | HA | A/Chicken/Netherlands/1/03 | Immune Technology | IT-003-008p |
| H7-13 | HA | A/chicken/Anhui/1/2013 | Sino Biological | 40103-V08H1 |
| H7-12 | HA | A/chicken/Jalisco/CPA1/2012 | Immune Technology | IT-003-0071p |
| H8-79 | HA | A/pintail duck/Alberta/114/1979 | Sino Biological | 11722-V08H1 |
| H9-99 | HA | A/Guinea fowl/Hong Kong/WF10/99 | Sino Biological | 11719-V08H1 |
| H9-97 | HA | A/chicken/Hong Kong/G9/97 | Sino Biological | 40036-V08H1 |
| H11-02 | HA | A/duck/Yangzhou/906/2002 | Immune Technology | IT-003-012p |
| H15-83 | HA | A/duck/Australia/341/1983 | Sino Biological | 11720-V08H1 |
| H16-99 | HA | A/black-headed gull/Sweden/5/99 | Sino Biological | 11711-V08H1 |

Nitrocellulose slides were treated with Blocker™ BLOTTO in TBS (Thermo Scientific™, Waltham, MA, USA) to prevent nonspecific binding. After blocking and washing the slides with PBS with 0.05% TWEEN® 20 (Merck-Millipore, Burlington, USA), plasma from bloodclots or serum was diluted 1:32 (v/v) in Blocker™ BLOTTO in TBS (Thermo Scientific™, Waltham, MA, USA) with 0.1% Tween™ 20 Surfact-Amps™ Detergent Solution (Thermo Scientific™, Waltham, MA, USA) and incubated for 1 hour at 37°C. After washing the slides, a 2-step detection was performed by diluting the conjugates 1:500 in Blocker BLOTTO with 0.1% Tween™ 20 Surfact-Amps and incubating for 1 hour at 37°C. First Goat anti-Canine IgG biotin (Thermo Scientific™, Waltham, MA, USA) was added to the fox and wolf samples and anti-ferret IgG biotin (Merck, Darmstadt, Germany) was added to the mustelid samples. After washing, secondary antibody mouse anti-biotin AlexaFluor 647 (Jackson Immuno Research, Ely, UK) was used for all samples. Fluorescence signal was measured using the Tecan power scanner (Männedorf Switzerland) and analyzed using ImaGene software. Further analysis and visualization was done using Graphpad PRISM (version 9.5.1). Significant differences in fluorescence signals were calculated using the Mann–Whitney U test.

### Haemagglutination inhibition (HI) assay

Serum and plasma samples were tested for the presence of H5-specific antibodies in HI assays according to the standard procedure, described previously [24]. All samples above cutoff in the PMA were screened for antibodies specific for LPAI H5 virus A/Mallard/Netherlands/96/2019 and clade 2.3.4.4b HPAI H5 virus A/Chicken/Netherlands/EMC-1/2018, inactivated with β-propiolactone (BPL). Fifty-three samples (2020-2022) which were below the cut-off for H5-HA1 in the PMA, of which 42 also did not bind to NP, were also tested in the HI assay and these samples showed no inhibition in the HI assay.

Serum/plasma samples were incubated for 16 hours at 37°C with *Vibrio cholerae* filtrate containing receptor destroying-enzyme (RDE) to remove non-specific inhibitors of haemagglutination activity, followed by a 1 hour incubation at 56°C. Subsequently, packed Turkey red blood cells (TRBC) were incubated for 1 hour at 4°C with treated serum/plasma to eliminate any additional non-specific haemagglutination activity that was not removed by RDE. Two-fold serial dilutions of the serum/plasma samples, starting at a dilution of 1:20 until 1:160, were prepared using phosphate-buffered saline (PBS) in U-bottomed 96 well microtitre plates. Serum/plasma dilutions were incubated with four haemagglutination units (HAU) of HPAI H5 virus A/Chicken/Netherlands/EMC-1/2018 (grown in Madin-Darby canine kidney cells) or LPAI H5 virus A/Mallard/Netherlands/96/2019 (grown in eggs) for 30 minutes at 37°C. A suspension of 1% Turkey red blood cells was added to the serum/plasma-virus dilutions. After incubation of 1 hour at 4°C, haemagglutination inhibition patterns were read. Negative controls, based on serum incubation without virus, were used to measure non-specific haemagglutination activity of each sample.

### Phylogeny and bioinformatics

All Dutch HPAI H5 sequences from the period 1-1-2020 to 14-2-2023 were downloaded from GISAID on 15 February 2023. Consensus sequences of viruses from all Dutch mammals, a random selection (https://www.randomizer.org/) of wild bird HPAI H5 sequences, and sequences from the Netherlands that were identical or most related (as determined by BLAST) were aligned using MUSCLE (3.8.425). A maximum likelihood phylogenetic tree was inferred using the IQ-TREE web server, using GTR+F+G4 as best fit model, with an approximate likelihood-ratio test (aLRT) as well as ultrafast bootstrapping with 1,000 replicates [32-34]. Flusurver was used to analyze carnivore sequences for mutations that indicate mammalian adaptation (http://flusurver.bii.a-star.edu.sg; Accessed 13 March 2023).

## Results

### Molecular detection of HPAI (H5) virus in wild carnivores

In total, 563 dead animals were submitted for HPAI viral RNA testing between 2020 and 2022. Of those, 174 animals were found dead and 389 were euthanized (Table 3). The majority of euthanized animals (n=375) were stone martens that had been trapped and euthanized in a meadow bird protection pilot (Supplemental materials). The remaining fourteen animals were euthanized following severe trauma or disease or as part of invasive animal control (e.g. the included raccoons). The total number of animals found dead did not differ significantly between seasons. Trapped stone martens were mainly collected in winter (53%) and spring (15%).

In total 20 out of 563 animals were HPAI H5 RNA positive. The highest incidence was found in foxes (9/31; 29%) and polecats (4/17; 24%; Table 2). Two positive foxes had been euthanized following neurological signs. Two out of four positive polecats showed neurological signs and were found dead later, whereas one ill polecat was euthanized with a likely parasitic infection. All remaining positive polecats and foxes were found dead, with the likely causes of death being trauma (n=6), infection (n=1) or unknown (n=1), based on macroscopic lesions. In addition, six trapped, apparently healthy, stone martens (6/400; 2%) were HPAI H5 RNA positive (Table 2). All positive foxes and polecats, as well as two of the six positive stone martens, were found in 2022 (Table 3). In addition, one positive badger was found dead in 2020, due to trauma, showing no signs of disease. Most positive animals were found in winter (75%; Table 3).

**Table 2.** Molecular and serological diagnostic results, wild carnivores, 2020-2022

|  | PCR | PMA |  | HI |
| --- | --- | --- | --- | --- |
|  | No H5 positive<br>/total no of<br>samples (%) | No NP reactive<br>/total no of<br>samples (%) | No H5* reactive<br>/total no of<br>samples (%) | HI<br>(A/Chicken/<br>Netherlands/<br>EMC-1/1/2018)<br>(%) |
| Pine marten (*Martes martes*) | 0/5 (0%) | 1/4 (25%) | 1/1 (100%) | N.T. |
| Polecat (*Mustela putorius*) | 4/17 (24%) | 2/11 (18%) | 2/2 (100%) | N.T. |
| Badger (*Meles meles*) | 1/51 (2%) | 4/48 (8) | 0/3 (0%) | N.T. |
| Stoat (*Mustela erminea*) | 0/4 (0%) | 1/4 (25%) | 0/1 (0%) | N.T. |
| Marten (*Martes spp.*) | 0/7 (0%) | 0/7 (0%) | N.T. | N.T. |
| Otter (*Lutra lutra*) | 0/7 (0%) | 0/6 (0%) | N.T. | N.T. |
| Stone marten (*Martes foina*) | 6/400 (2%) | 135/269 (50%) | 71/126 (56%) | 19/65 (29%) |
| Fox (*Vulpes vulpes*) | 9/31 (29%) | 19/27 (70%) | 7/19 (37%) | 2/6 (33%) |
| Raccoon (*Procyon lotor*) | 0/8 (0%) | 0/8 (0%) | N.T. | N.T. |
| Weasel (*Mustela nivalis*) | 0/24 (0%) | 0/14 (0%) | N.T. | N.T. |
| Wolf (*Canis lupus*) | 0/9 (0%) | 3/7 (43%) | 0/3 (0%) | N.T. |

\*clade 2.3.4.4

**Table 3.** General Characteristics of the Wild Carnivores with and without AIV (H5) Infection

|  |  | H5 positive | H5 negative |
| --- | --- | --- | --- |
| **Total** | **Total (563)** | **20 (100%)** | **543 (100%)** |
| Season | Winter (251) | 15 (75%) | 236 (43%) |
|  | Spring (104) | 2 (10%) | 102 (19%) |
|  | Summer (47) | 3 (15%) | 44 (8%) |
|  | Autumn (69) | 0 (0%) | 69 (13%) |
|  | Unknown (92) | 0 (0%) | 92 (17%) |
| Year | 2020 (132) | 1 (5%) | 131 (24%) |
|  | 2021 (279) | 4 (20%) | 275 (51%) |
|  | 2022 (152) | 15 (75%) | 137 (25%) |
| Status | Found dead (174) | 11 (55%) | 163 (30%) |
|  | Euthanized (389) | 9 (45%) | 380 (70%) |
| Possible cause of death | Trauma * (504) | 13 (65%) | 491 (90.5%) |
|  | Infection (34) | 6 (30%) | 28 (5%) |
|  | Neoplasia (2) | 0 (0%) | 2 (0.5%) |
|  | Other (5) | 0 (0%) | 5 (1%) |
|  | Unknown (18) | 1 (5%) | 17 (3%) |

\* Including trapped stone martens

Of A(H5) positive animals, 12 out of 20 throat swabs tested positive for H5 RNA, as well as 5 out of 19 available lung samples and 6 out of 16 brain samples. Eight out of 18 available rectal swabs were A(H5) positive (supplemental materials).

A total of seven (almost) complete HPAI H5 virus genome sequences from infected animals were obtained: five from foxes, one from a polecat and one from a stone marten. Of these, three sequences from foxes (3/7; 43%) contained the E627K substitution in the PB2 open reading frame. Substitution T271A was found in the PB2 open reading frame of the virus sequence from the polecat. This substitution was also found in the infected mink reported recently in Spain[13]. These PB2 substitutions are known to be associated with increased replication in mammalian cells. All sequences obtained from the HPAI H5 infected wild carnivores in this study cluster phylogenetically with sequences obtained from birds from the Netherlands (Figure 2). Two fox HA sequences, as well as the stone marten sequence and another Dutch fox sequence (outside this study) cluster together, as well as with multiple bird sequences, but these carcasses were not found in the same timeframe or location.

**Figure 1A.**
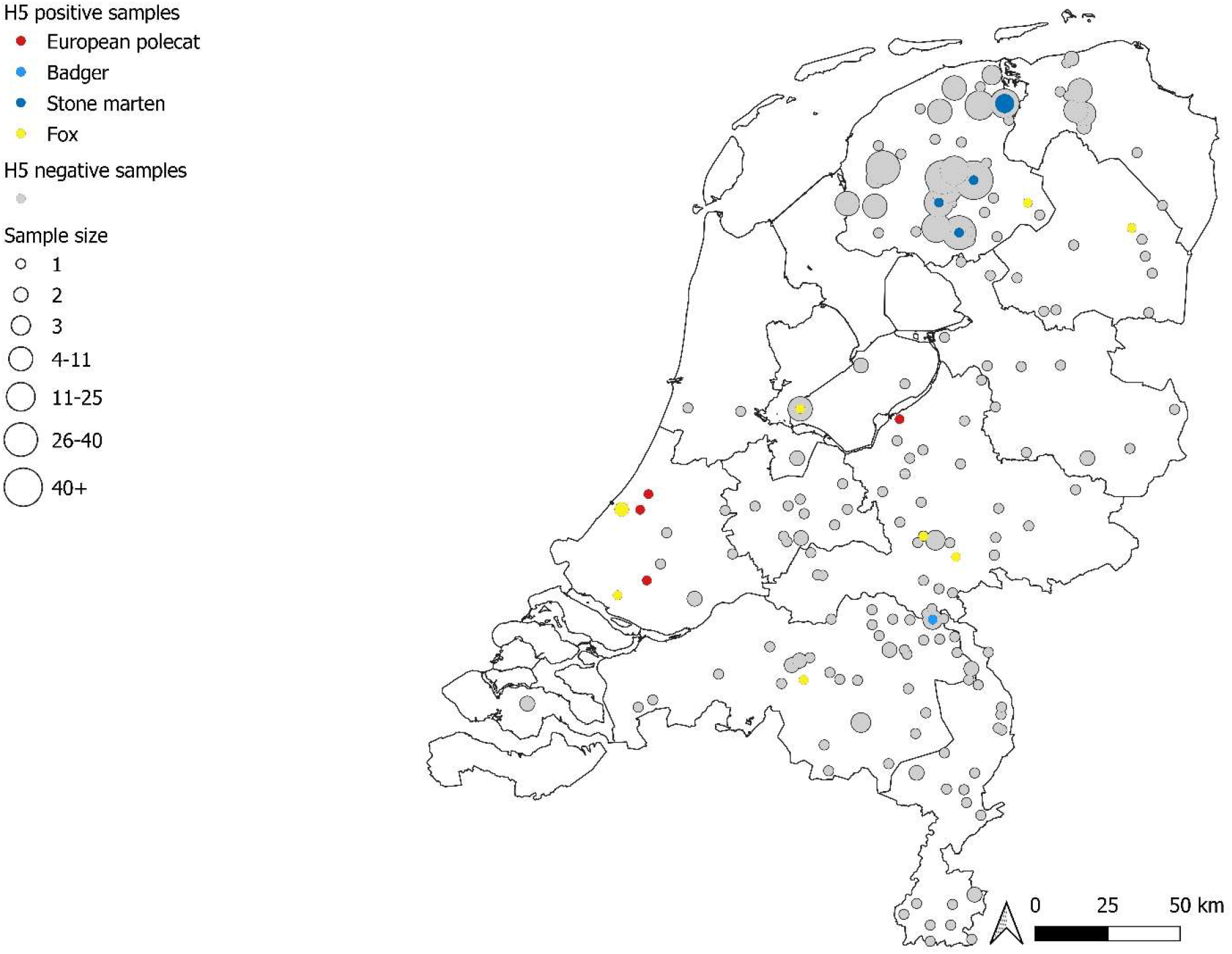
**HPAI H5 PCR positive wild carnivores, 2020-2022**

**Figure 1B.**
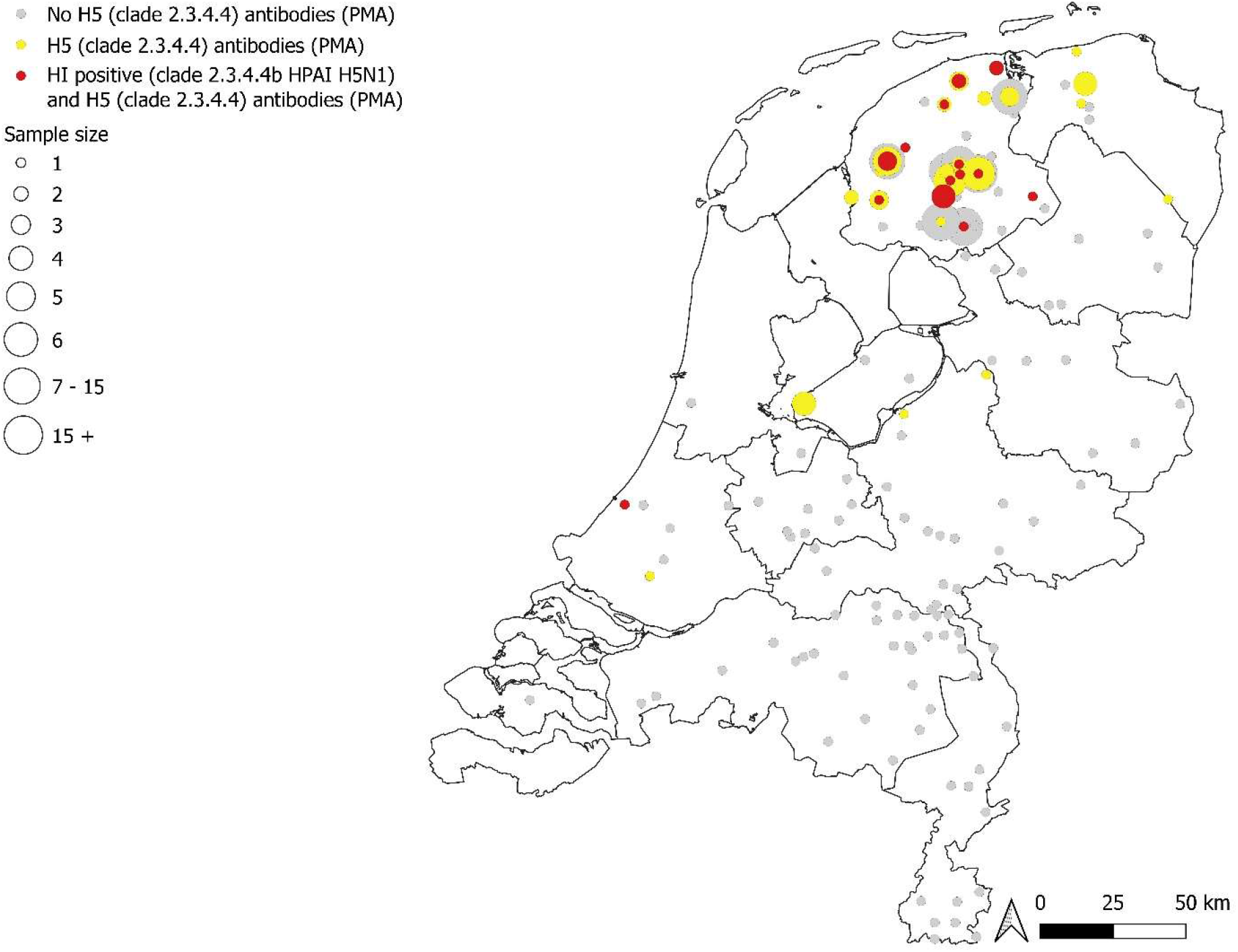
**HPAI H5 (clade 2.3.4.4b) antibody positive wild carnivores, 2020-2022**

**Figure 2.**
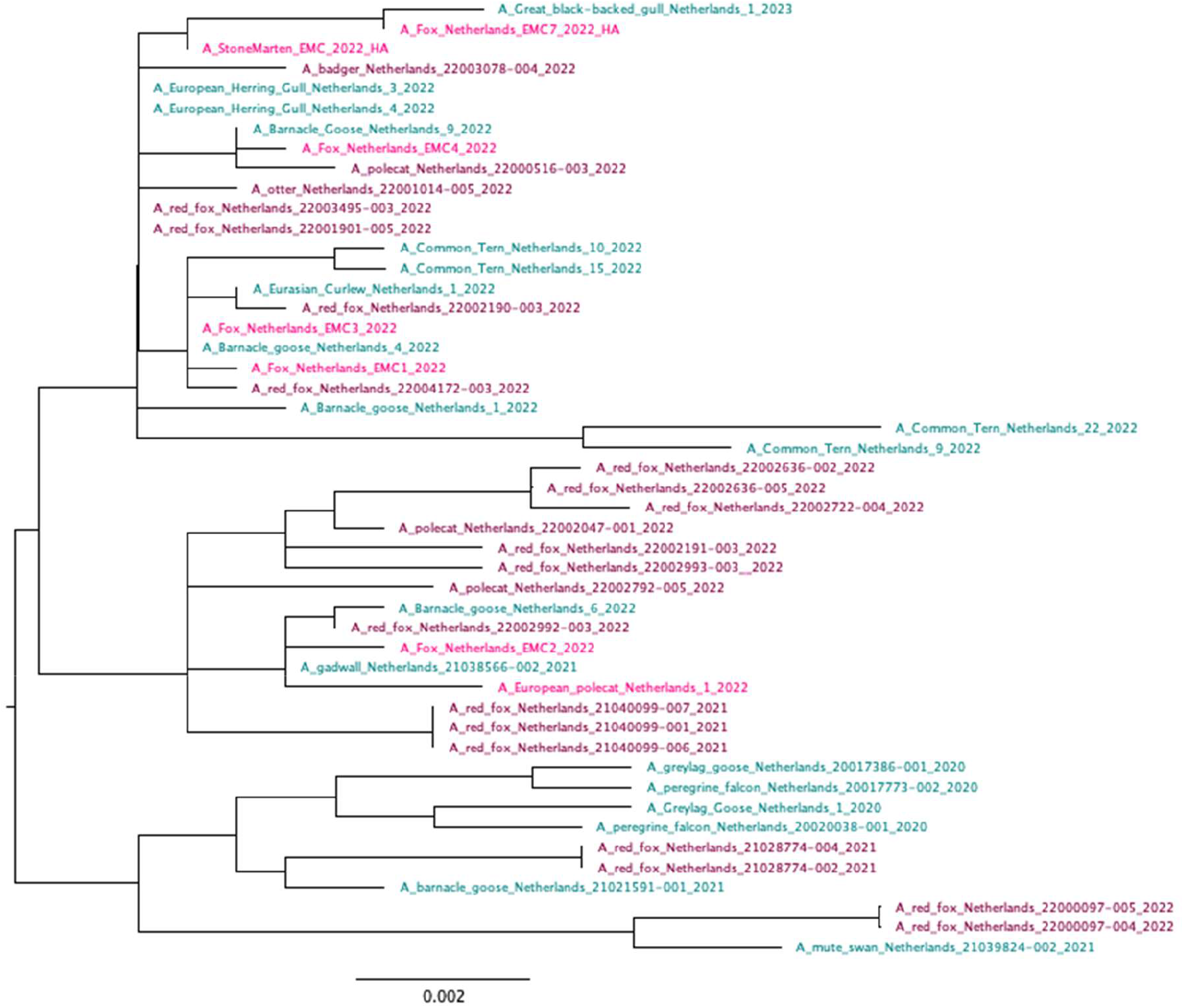
Phylogenetic tree for the HA gene of Dutch carnivore HPAI H5 virus sequences. Phylogenetic tree of all Dutch HPAI H5 virus mammal sequences (Sequences from this study in magenta, other sequences from GISAID in prune) and a selection of A(H5) sequences obtained from infected wild birds (colored in teal), 2020-2022

### Serological analyses

In total, samples from 405 dead wild carnivores, from the period 2020-2022, were screened with the PMA. In 165 wild carnivores (165/405; 41%) Influenza A NP-binding antibodies were detected (Table 2; Figure 3). There was a significant yearly increase of the number of animals, sampled between 2020 and 2022, that showed NP reactivity (P<0.001; Figure 3). In total, 20% (81/405) of all tested carnivores showed binding of the clade 2.3.4.4 H5-HA1 antigen in the PMA (Table 2; Figure 3). Of those, 21 (21/71; 30%) samples showed inhibition in the HI assay against the HPAI H5 clade 2.3.4.4b virus A/Chicken/Netherlands/EMC-1/2018, with titers ranging from 20 to ≥ 160. None of the plasma and serum samples showed inhibition to the LPAI H5 virus A/Mallard/Netherlands/96/2019. Consistent with the PCR results, a significant portion of foxes and stone martens had antibodies against HPAI H5 viruses as measured with both PMA and HI assay.

**Figure 3.**
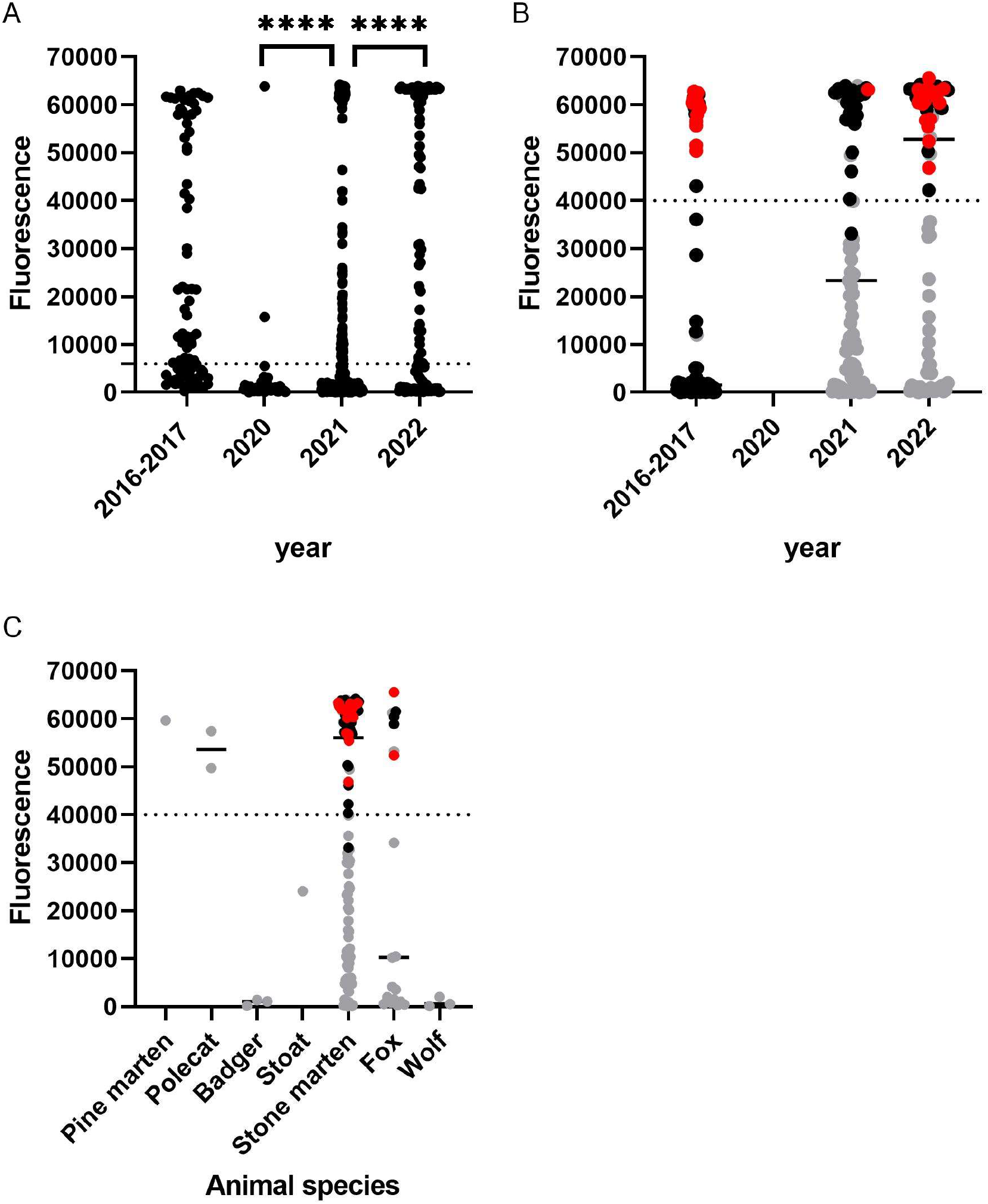
H5 serology in wild carnivores, 2016-2017; 2020-2022. A. Influenza A NP serology results by year, data points show H7 NP signals. When H1 NP exceeded the cut-off (PFU 6000), but H7 NP did not, we show H7 NP fluorescent signals (n=4). B. H5 (clade 2.3.4.4) serology results by year. Only sera above the cutoff for NP antigen (PFU 6000) were included. Data points in grey were tested in PMA but negative, black data points were H5 positive in PMA and negative in HI assay and red dots were positive in both assays. C. H5 (clade 2.3.4.4) serology results by carnivore species. Only sera above the cutoff for NP antigen (PFU 6000) were included. Data points in grey were tested in PMA but negative, black data points were H5 positive in PMA and negative in HI assay and red dots were positive in both assays.

In addition, 73 fox sera collected by hunters, as well as plasma from one pole cat and three stone martens (obtained from dead animals) from 2016-2017, collected in the Netherlands, were tested with the PMA and HI assay. NP binding antibodies were detected in 50 (50/77; 65%) samples (1 stone marten and 49 foxes). HA1-H5 binding antibodies were detected in 23 sera (23/77; 30%), all from foxes. Eleven foxes were also positive for HPAI H5 clade 2.3.4.4b antibodies in the HI assay, with HI titers ranging from 20-80 but negative against the LPAI H5 virus A/Mallard/Netherlands/96/2019. The other sera were negative in both assays.

## Discussion

This study shows that the exposure and number of HPAI H5 virus infections between 2020-2022 in wild carnivores was much higher than previous reports suggested. Our serology studies demonstrated that significant proportions of foxes and stone martens that were found dead had antibodies against HPAI H5 virus. The same animal species also tested positive for HPAI H5, with 29% of dead foxes and 24% of dead polecats testing RNA positive. Only four out of 20 H5 positive carnivores showed neurological complications. Therefore, carnivores that do not show abnormal neurological behavior or encephalitis may also be infected frequently, and serology indicated that exposure does not always lead to mortality. Data from 2016 and 2017 indicated that wild carnivore infections were also occurring during previous outbreaks of clade 2.3.4.4b HPAI H5 viruses.

The current HPAI H5 virus outbreak is the largest outbreak ever reported, and since 2021, HPAI H5 viruses can be detected year-round in Europe. Since most wild carnivore infections are likely caused by predation or direct contact with sick or dead wild birds [8,9], both the magnitude and timing of infections in birds will affect the number of carnivore infections that occur. Our study indeed suggests an increase in wild carnivore infections in 2022, versus 2020 and 2021, based on molecular screening and serology. However, the nature of the survey precludes robust conclusions. The results of the serological analysis of sera from 2016 and 2017 suggest that exposure rates may also have been high during previous clade 2.3.4.4b HPAI H5 virus outbreaks. The drop in seroprevalence between 2017 and 2020 is likely explained by antibody waning and lack of exposure of young animals in the absence of HPAI H5 virus outbreaks in the Netherlands between 2018-2020.

Notably, the HPAI H5 virus infected stone martens in this study were trapped, not found dead, suggesting that HPAI virus infection does not always result in noticeable or fatal disease in mammals. Also, the observed numbers of foxes, polecats and stone martens with antibodies against HPAI H5 virus with trauma as the most likely cause of death, suggests that infections may not always lead to severe disease, although trauma may happen more frequently for sick animals. This observation is different from the currently available body of evidence, describing ante mortem neurological signs in most of the cases or positive animals that were found dead [35,36]. However recent ferret laboratory infections with clade 2.3.4.4b HPAI H5 viruses also confirm the possibility of mild disease following infection in mustelids [37]. Furthermore, in the HPAI H5 virus infected farmed mink in Spain, notable lesions in the lungs, but not brains, were most notable [13]. Therefore, infections with clade 2.3.4.4b HPAI H5 viruses do not always result in neuropathy or death, and disease presentation may even differ between different viruses within clade 2.3.4.4b, complicating surveillance and risk assessment [16].

In agreement with previous studies [8,38] our sequence data shows repeated emergence of mammalian adaptation markers in the carnivores in this study. The E627K amino acid substitution in the PB2 open reading frame is known to be a molecular determinant of host range [39] and is an important virulence factor in HPAI H5N1 human infections [40]. Although zoonotic infections with HPAI H5 viruses have been reported, the number of human infections with the currently circulating clade 2.3.4.4b viruses in Europe and the Americas is limited, and most patients did not exhibit severe symptoms [36]. Two exceptions are the recently reported cases of a nine-year-old girl, admitted to an intensive care unit (ICU) in Ecuador and a 53-year-old man from Chile, also admitted to the ICU with dyspnea and respiratory distress (https://www.who.int/emergencies/disease-outbreak-news/item/2023-DON434; https://www.paho.org/en/documents/briefing-note-human-infection-caused-avian-influenza-ah5-virus-chile-march-31-2023 retrieved on 11-4-2023). However, with the current size of the outbreak and seeming increasing number of mammal infections, opportunities for the virus to adapt from avian to mammalian hosts are increasing. This may increase the risk of acquisition of properties for efficient mammal-to-mammal transmission, resulting in significant risks for public health [16,41].

The role of wild carnivores in the transmission and spread of HPAI H5 virus is unclear. Possible mammal-to-mammal transmission has occurred during HPAI H5 outbreaks in seals in the USA [14] and in South American sea lions (*Otaria flavescens)* in Peru [15]. Furthermore, during the recent outbreak in farmed mink [13], a species closely related to marten species, transmission of HPAI H5N1 virus within the farm was likely. This may also occur in wild mustelids and other carnivores, although this has not been described to date [8]. The viral genome sequences in this study mainly clustered with virus sequences from birds, and there was only very limited geographical and/or temporal clustering between the positive wild carnivores. Therefore, individual infections after exposure to infected birds, is the most likely infection route. Moreover, many carnivores have a solitary life style and rarely come into contact with other carnivores, even within their own species. Possible spill-back of viruses from carnivores to wild birds has also been suggested, for example after predation of infected carcasses by birds of prey [16]. Introductions of viruses with amino acid changes that facilitate replication in mammals into wild birds would greatly increase chances of further spread.

The interactions between wild carnivores and domestic animals are of concern, because HPAI H5 virus infections in domestic animals, particularly farmed animals, can lead to severe disease, mortality and high costs, as well as risks to public health due to increased risk of mammalian adaptation. Moreover, there is a risk of reassortment in mammalian species susceptible to both human and avian viruses, such as pigs and mustelids [42,43]. Previous research of SARS-CoV-2 and Canine Distemper Virus (CDV) on mink farms has shown that wild carnivores frequently enter farms and introduce novel viruses to the farmed mink [44,45]. Similar to wild carnivores, domestic carnivores also catch and eat birds, and can get exposed to avian influenza viruses. This was exemplified by a recent HPAI H5 virus infection in a cat (https://wahis.woah.org/#/in-review/4807; Avian influenza virus infects a cat | Anses -Agence nationale de sécurité sanitaire de l’alimentation, de l’environnement et du travail retrieved on 11-4-2023). Although domestic carnivores are usually kept in small numbers, they are in close contact with humans, with a risk of animal-to-human transmission of adapted HPAI viruses. Furthermore, avian influenza infections in rodents have been described [46], but so far rarely any monitoring or field research is targeting them.

## Conclusion

Here, we show that foxes, polecats and stone martens are infected frequently with clade 2.3.4.4b HPAI H5 viruses, with and without clear neurological signs or mortality. Some of the viruses showed evidence for adaptation to mammals. This demonstrates the need for increased surveillance of all wild carnivores to monitor infections and mutations, irrespective of neurological signs. Increased surveillance should also include other wild and domesticated animal species, such as swine, mink, cats, dogs and more. Also, avian influenza infections in rodents have been described, but so far rarely any monitoring or field research is targeting them. Especially species that are susceptible to human as well as avian influenza viruses are of relevance, because of the risk of reassortment. Initiating such active monitoring of both live and deceased animals will not only contribute to our understanding of the epidemiology and pathogenesis of currently circulating HPAI virus strains, but will also aid in the timely identification of novel high-risk variants.

## Acknowledgements

We kindly thank the members of the DWHC and VPDC from the Utrecht University for their technical assistance. Special thanks to Ruby Wagensveld and Natashja Ennen-Buijs for their administrative and technical contribution. We thank the volunteers for their assistance during the necropsy sessions. Marjan Boter, Merve Tok, Aurora Horstink and Charety Koense are thanked for their assistance in sample processing. We thank Felicity Chandler for her help with printing and testing the PMA slides. We gratefully acknowledge all data contributors, i.e., the Authors and their Originating laboratories responsible for obtaining the specimens, and their Submitting laboratories for generating the genetic sequence and metadata and sharing via the GISAID Initiative.

