## Supplemental Table 1 for "High number of HPAI H5 Virus Infections and Antibodies in Wild Carnivores in the Netherlands, 2020-2022"

### Supplemental Table 1. RT-PCR (H5 screening) results per sample type, wild carnivores, 2020-2022

|  | **Cycle threshold per material type** | | | | |
| --- | --- | --- | --- | --- | --- |
| **Animal** | **Throat swab** | **Colon swab** | **Lung** | **Brain** | **Feces** |
| Pole cat | **36** | Not detected | Not detected | Not detected | No feces |
| Pole cat | **18** | **23** | **19** | **18** | No feces |
| Pole cat | Not detected | Not tested | Not detected | **40** | No feces |
| Pole cat | **30** | **31** | **30** | Not detected | No feces |
| Badger | Not tested | Not tested | Not detected | **34** | Not detected |
| Stone marten | **36** | Not detected | Not detected | Not detected | No feces |
| Stone marten | **36** | **28** | Not detected | Not detected | No feces |
| Stone marten | Not detected | **34** | Not detected | Not detected | No feces |
| Stone marten | **28** | Not detected | Not detected | Not detected | **36** |
| Stone marten | **35** | Not detected | Not detected | Not detected | Not tested |
| Stone marten | Not tested | **35** | Not tested | Not detected | No feces |
| Fox | **26** | **26** | **22** | **26** | No feces |
| Fox | **30** | Not tested | Not detected | Not detected | No feces |
| Fox | **27** | **29** | No lung | No brain | No feces |
| Fox | Not tested | **23** | **20** | No brain | No feces |
| Fox | Not tested | Not tested | **32** | No brain | No feces |
| Fox | **38** | Not tested | Not tested | No brain | No feces |
| Fox | Not tested | Not tested | Not tested | **25** | No feces |
| Fox | **35** | Not tested | Not detected | **22** | No feces |
| Fox | Not detected | Not tested | Not detected | Not detected | No feces |
