## Supplemental Table 2 for "High number of HPAI H5 Virus Infections and Antibodies in Wild Carnivores in the Netherlands, 2020-2022"

### Supplemental Table 2. Characteristics H5 positive wild carnivores, 2020-2022

| **Badger** | **Total (51)** | **1 (100%)** | **50 (100%)** |
| --- | --- | --- | --- |
| Season | Winter (7) Spring (15) Summer (10) Autumn (19) | 0 (0%) 0 (0%) 1 (100%) 0 (0%) | 7 (14%) 15 (30%) 9 (18%) 19 (38%) |
| Year | 2020 (23) 2021 (21) 2022 (7) | 1 (100%) 0 (0%) 0 (0%) | 22 (44%) 21 (42%) 7 (14%) |
| Status | Found dead (48) Euthanized (3) | 1 (100%) 0 (0%) | 47 (94%) 3 (6%) |
| Possible cause of death | Trauma (41) Infection (4) Neoplasia (1) Other (2) Unknown (3) | 1 (100%) 0 (0%) 0 (0%) 0 (0%) 0 (0%) | 40 (80%) 4 (8%) 1 (2%) 2 (4%) 3 (6%) |
| **Fox** | **Total (31)** | **9 (100%)** | **22 (100%)** |
| Season | Winter (16) Spring (8) Summer (4) Autumn (3) | 5 (56%) 2 (22%) 2 (22%) 0 (0%) | 11 (50%) 6 (27%) 2 (9%) 3 (14%) |
| Year | 2020 (2) 2021 (5) 2022 (24) | 0 (0%) 0 (0%) 9 (100%) | 2 (9%) 5 (23%) 15 (68%) |
| Status | Found dead (26) Euthanized (5) | 7 (78%) 2 (22%) | 19 (86%) 3 (14%) |
| Possible cause of death | Trauma (12) Infection (13) Unknown (6) | 5 (56%) 3 (33%) 1 (11%) | 7 (32%) 10 (45%) 5 (23%) |
| **Pole cat** | **Total (17)** | **4 (100%)** | **13 (100%)** |
| Season | Winter (6) Spring (2) Summer (6) Autumn (3) | 4 (100%) 0 (0%) 0 (0%) 0 (0%) | 2 (15.5%) 2 (15.5%) 6 (46%) 3 (23%) |
| Year found | 2020 (2) 2021 (5) 2022 (10) | 0 (0%) 0 (0%) 4 (100%) | 2 (15.5%) 5 (38.5%) 6 (46%) |
| Status | Found dead (16) Euthanized (1) | 3 (75%) 1 (25%) | 13 (100%) 0 (0%) |
| Possible cause of death | Trauma (9) Infection (5) Unknown (3) | 1 (25%) 3 (75%) 0 (0%) | 8 (62%) 2 (15%) 3 (23%) |
| **Stone marten (found dead)** | **Total (25)** | **0 (100%)** | **25 (100%)** |
| Season | Winter (7) Spring (8) Summer (2) Autumn (8) | 0 (0%) 0 (0%) 0 (0%) 0 (0%) | 7 (28%) 8 (32%) 2 (8%) 8 (32%) |
| Year found | 2020 (5) 2021 (12) 2022 (8) | 0 (0%) 0 (0%) 0 (0%) | 5 (20%) 12 (48%) 8 (32%) |
| Status | Found dead (25) Euthanized (0) | 0 (0%) 0 (0%) | 25 (100%) 0 (0%) |
| Possible cause of death | Trauma (12) Infection (8) Other (3) Unknown (2) | 0 (0%) 0 (0%) 0 (0%) 0 (0%) | 12 (48%) 8 (32%) 3 (12%) 2 (8%) |
| **Stone marten (trapped)** | **Total (375)** | **6 (100%)** | **369 (100%)** |
| Season | Winter (202) Spring (54) Summer (6) Autumn (21) Unknown (92) | 6 (100%) 0 (0%) 0 (0%) 0 (0%) 0 (0%) | 196 (53%) 54 (14.5%) 6 (1.5%) 21 (6%) 92 (25%) |
| Year | 2020 (78) 2021 (210) 2022 (87) | 0 (0%) 4 (67%) 2 (33%) | 78 (21%) 206 (56%) 85 (23%) |
| Status | Found dead (0) Euthanized (375) | 0 (0%) 6 (100%) | 0 (0%) 369 (100%) |
| Possible cause of death | Trauma (375) | 6 (100% | 369 (100%) |
| **H5 negative wild carnivores** | **Total (64)** | **0 (100%)** | **64 (100%)** |
| Season | Winter (13) Spring (17) Summer (19) Autumn (15) | 0 (0%) 0 (0%) 0 (0%) 0 (0%) | 13 (20%) 17 (27%) 19 (30%) 15 (23%) |
| Year found | 2020 (34) 2021 (41) 2022 (25) | 0 (0%) 0 (0%) 0 (0%) | 22 (34%) 26 (41%) 16 (25%) |
| Status | Found dead (59) Euthanized (5) | 0 (0%) 0 (0%) | 59 (92%) 5 (8%) |
| Possible cause of death | Trauma (55) Infection (4) Neoplasia (1) Unknown (4) | 0 (0%) 0 (0%) 0 (0%) 0 (0%) | 55 (86%) 4 (6%) 1 (2%) 4 (6%) |
