## Supplementary figures and images for "High number of HPAI H5 Virus Infections and Antibodies in Wild Carnivores in the Netherlands, 2020-2022"

### Supplemental Figure 1

#### Supplemental Figure 1. HPAI H5 (clade 2.3.4.4b) antibody positive wild carnivores, per year

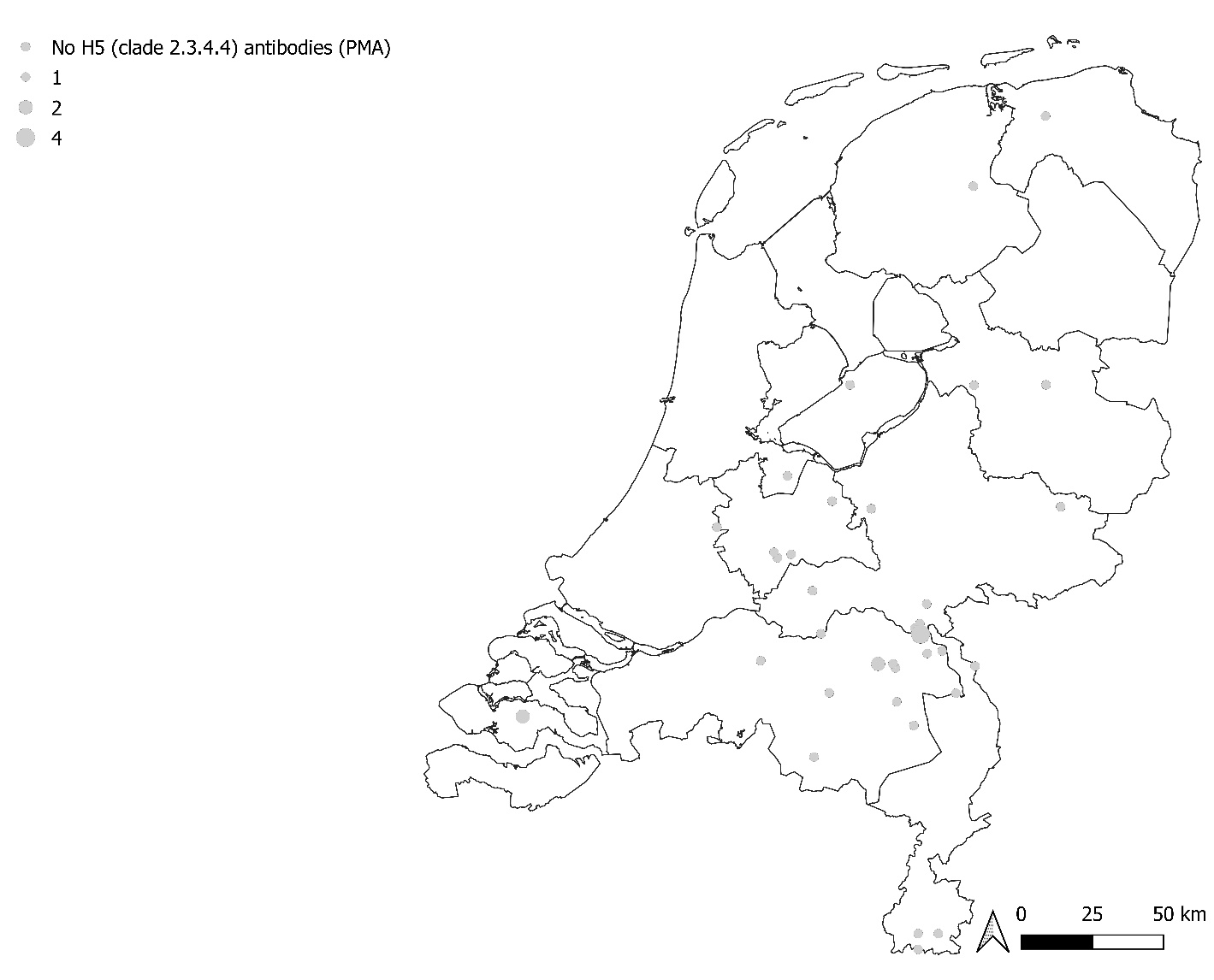

1A. 2020

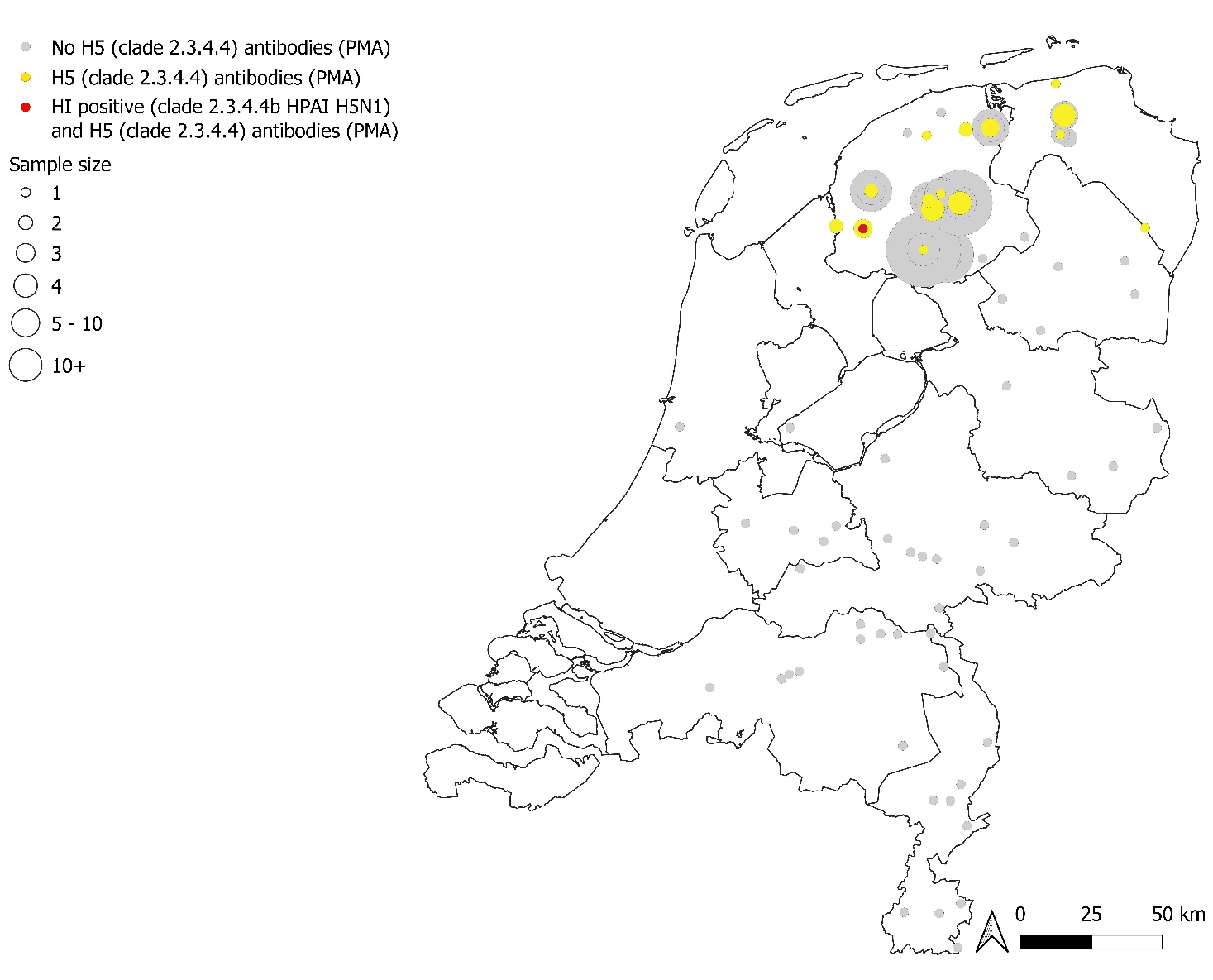

1B. 2021

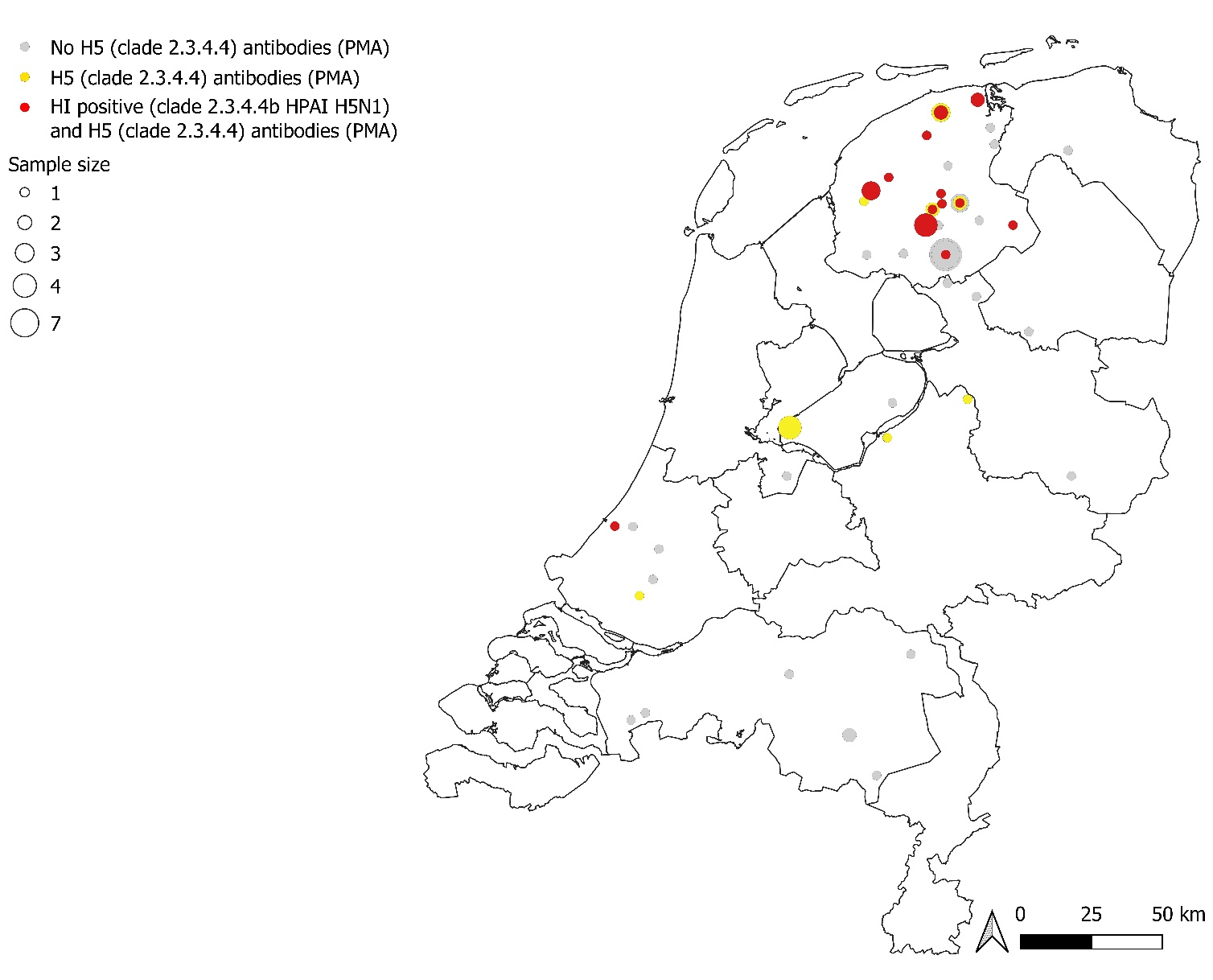

1C. 2022
